# An improved workflow for rapid, large-scale protein production in HEK293 cells via antibiotic enrichment after lentiviral transduction

**DOI:** 10.64898/2026.03.07.710266

**Authors:** Ester Behiels, Anushka Nair, Adrien Doridant, Jonathan Elegheert

## Abstract

Lentiviral transduction of HEK293-derived expression cells provides a robust and scalable approach for large-scale protein production for structural and biochemical studies. Building on our previously reported platform, we introduce an improved workflow that decouples cell enrichment from target protein expression by enabling constitutive antibiotic selection of transduced cells prior to induction. The key advance is the use of orthogonal antibiotic-resistance cassettes to stringently enrich transduced cells, eliminate non-transduced cells, improve population homogeneity, and enable multi-vector co-selection for heteromeric assemblies and complexes. We provide two complementary transfer-vector suites. pHR-AB-CMV-TetO_2_ delivers maximal expression and supports inducible control in TetR-expressing lines while driving strong constitutive expression in non-TetR lines. pHR-AIO-AB (“all-in-one”) encodes the transactivator, resistance marker, and gene of interest on a single construct to enable tightly controlled doxycycline-inducible expression in standard HEK293 lines, and is readily adaptable to other mammalian cell types. Both suites are available with puromycin, blasticidin, hygromycin, or zeocin markers, enabling straightforward co-infection and orthogonal multi-antibiotic selection of stable populations expressing multiple transgenes. They are well suited to demanding targets such as membrane proteins and multi-subunit assemblies. The protocol details the step-by-step generation of highly enriched, inducible HEK293 populations within 3–4 weeks.

## Introduction

### Development of the protocol

Producing milligram quantities of correctly folded and post-translationally modified proteins is essential for structural and biochemical studies. Mammalian expression systems, especially HEK293 (human embryonic kidney 293)-derived cell lines, are often the most reliable option for challenging eukaryotic targets such as membrane proteins, large multi-domain proteins, and multi-subunit assemblies. Lentiviral transduction of mammalian expression cells has emerged as a powerful tool for generating stable expression cell lines suitable for structural studies ^1^ because the vectors integrate into the host genome and support durable, uniform expression.

Our previously described lentiviral expression system ^1^ showed that polyclonal stable cell lines could be generated rapidly without the need for clonal selection, enabling time- and cost-efficient protein production. However, two practical limitations remained. First, there was no independent antibiotic (AB) selection to enrich transduced cells prior to transgene induction. Second, streamlined co-expression of multiple proteins was not supported, thereby complicating studies that require heteromeric complexes, receptor–ligand pairs, or engineered antibody formats.

Here we present an updated lentiviral platform that addresses both issues while preserving the simplicity and scalability of the original method. The key advance is antibiotic enrichment of transduced populations prior to induction, producing homogeneous, high-expressing polyclonal lines. Orthogonal resistance cassettes enable co-infection and multi-antibiotic co-selection of expression cells, facilitating coordinated assembly of complexes and heteromeric assemblies. The protocol remains compatible with common HEK293 variants and supports rapid generation of stable, inducible production lines suitable for demanding structural biology applications.

### Overview of the procedure

The workflow mirrors a standard two-step lentiviral transduction ^1^ but adds an antibiotic-enrichment phase enabled by our redesigned transfer vectors (Fig. 1a). Briefly, a dedicated HEK293T producer line (Lenti-X ΔPKR) is transiently co-transfected with the transfer plasmid (either pHR-AB-CMV-TetO_2_ or pHR-AIO-AB), the packaging plasmid (psPAX2) and envelope plasmid (pMD2.G) to generate lentiviral particles. The clarified supernatant is then used, typically without concentration, to transduce HEK293-derived expression cells. Because the transfer vectors constitutively express an antibiotic resistance cassette, transduced cells can be stringently enriched by antibiotic treatment before any induction of the gene of interest (GOI). The result is a polyclonal, stable HEK293 population that is ready for scale-up and protein production.

**Fig. 1.**
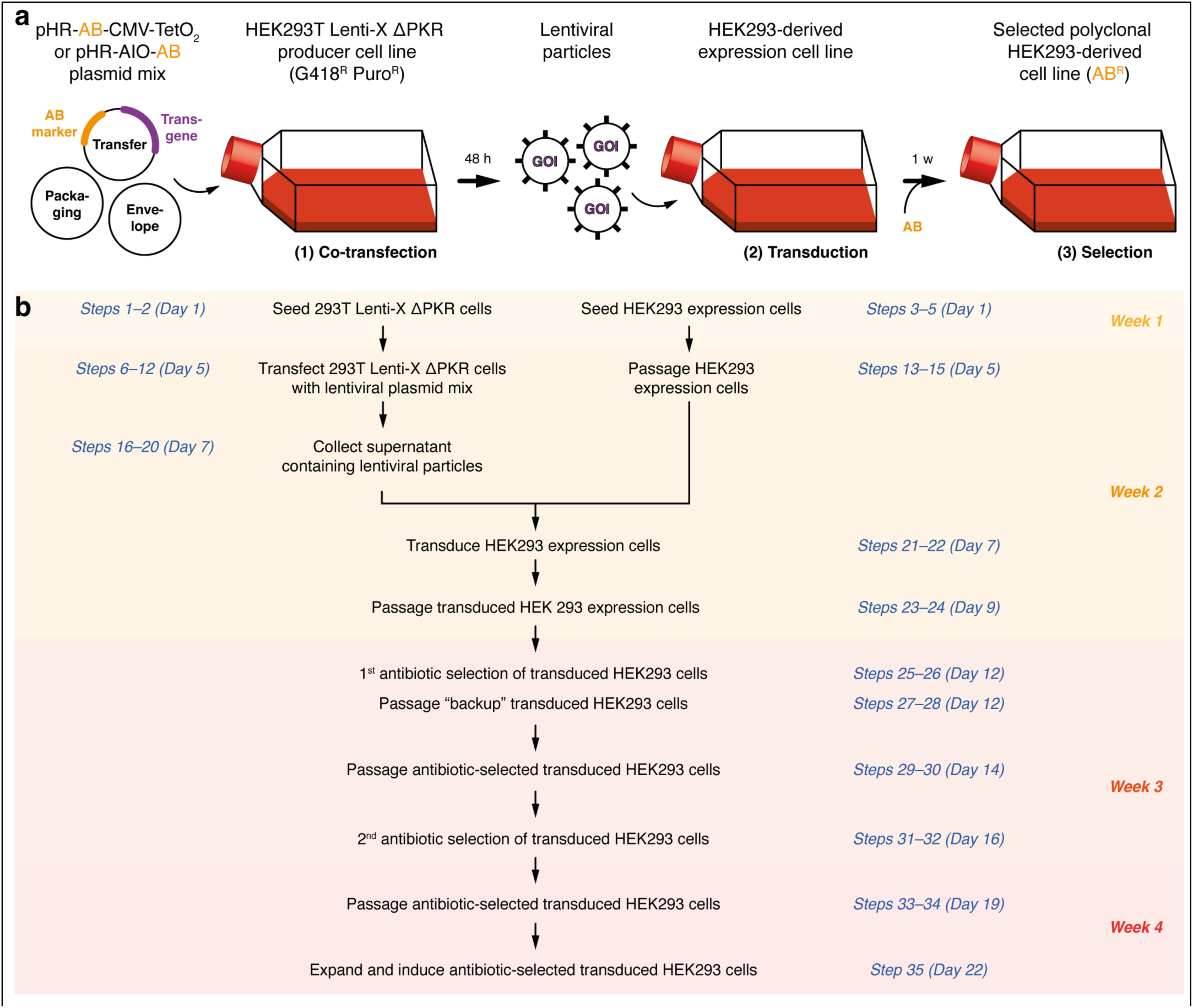
Principle and workflow of lentiviral infection and antibiotic enrichment. **a**, Schematic of the basic procedure: (1) the HEK293T Lenti-X ΔPKR producer line is co-transfected with transfer plasmid (pHR-AB-CMV-TetO_2_ or pHR-AIO-AB; carrying the gene of interest), packaging plasmid (psPAX2) and envelope plasmid (pMD2.G) to generate lentiviral particles; (2) HEK293-derived expression cells are transduced and (3) transduced cells are enriched by antibiotic selection. **b,** Flowchart of the protocol with indicative timings. Week numbering corresponds to a procedure initiated on a Thursday afternoon for convenience (Day 1). Abbreviations: AB, antibiotic; CMV, (human) cytomegalovirus; TetO_2_, two tandem tetracycline operator sites; HEK293, human embryonic kidney 293; PKR, (host) protein kinase R; G418^R^, geneticin resistance; Puro^R^, puromycin resistance.

The procedure is organised into four stages (Fig. 1b). Stage 1 (Steps 1–20) – Virus production: transient co-transfection of HEK293T Lenti-X ΔPKR cells with transfer, packaging and envelope plasmids, followed by harvest of lentiviral supernatant after 48 h. Stage 2 (Steps 21–24) – Transduction: filtration of the supernatant, addition of polybrene, infection of HEK293 expression cells (adherent HEK293T or suspension-adapted HEK293S GnTI^−^ variants) and passaging of the transduced cells. Stage 3 (Steps 25–34) – Antibiotic enrichment: repeated cycles of antibiotic selection (puromycin, blasticidin, hygromycin and/or zeocin) to enrich the stably transduced population to >95%. Stage 4 (Step 35) – Large-scale protein production (as detailed previously ^1^): expansion of the fully selected population (in adherent or suspension formats) and either induction of expression with doxycycline (Dox) or constitutive expression, depending on the chosen combination of vector and cell line. Although Fig. 1 illustrates transduction with a single GOI, the same workflow is applied to co-infection with multiple lentiviral preparations carrying orthogonal antibiotic markers, enabling stable co-expression of multi-subunit complexes.

### Features of the pHR-AB-CMV-TetO_2_ transfer plasmids

The updated design of this lentiviral plasmid suite was made to address some shortcomings of the original pHR-CMV-TetO_2_ system ^1^, where both the GOI and the selection marker gene (coding for a fluorescent protein or an antibiotic resistance marker) were under control of the same promoter (the CMV-MIE promoter), necessitating expression of the GOI to perform selection of transduced cells. Our goal was thus to redesign the system to include a separate, independent promoter for the constitutive expression of an antibiotic resistance marker. This new vector configuration, termed pHR-AB-CMV-TetO_2_ (Fig. 2a, Table 1 and Table 2), thus allows for independent expression of the GOI, with induced expression only performed after antibiotic selection, enrichment and scale-up of the cell line.

**Fig. 2.**
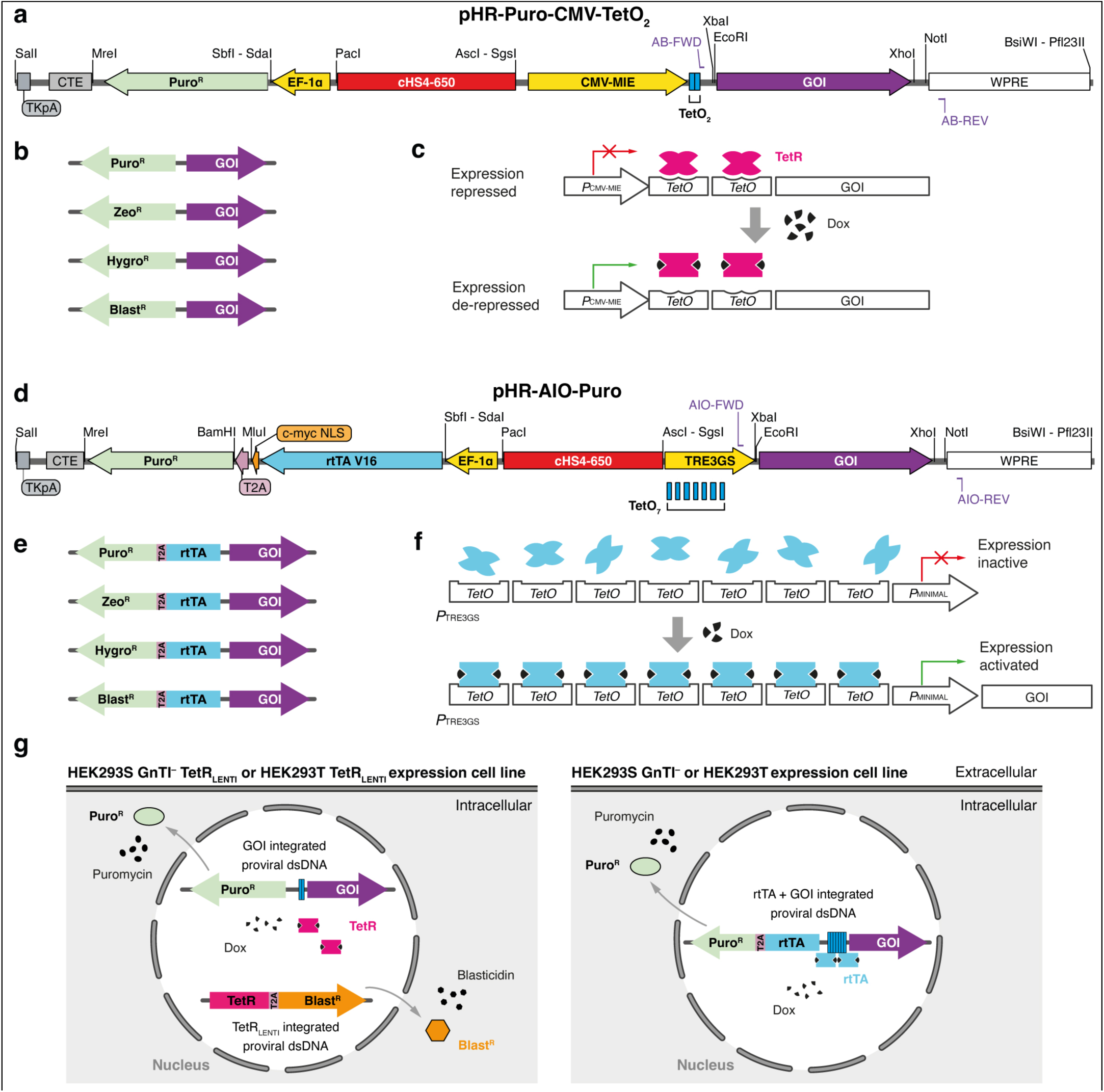
Design and features of the pHR-AB-CMV-TetO_2_ and pHR-AIO-AB lentiviral transfer vectors. **a**, Vector architecture of pHR-AB-CMV-TetO_2_ (Puro^R^ variant). The CMV-MIE promoter drives GOI expression, whereas a compact EF-1*α* promoter independently drives expression of the antibiotic-resistance cassette to enable stringent enrichment of transduced cells before induction. All genetic elements are flanked by unique restriction sites. **b**, Antibiotic variants of the pHR-AB-CMV-TetO_2_ system, available with puromycin, blasticidin, hygromycin or zeocin cassettes, permitting single-vector selection or orthogonal co-selection when co-infecting with multiple constructs. **c**, Inducible expression mechanism of the pHR-AB-CMV-TetO_2_ vectors. In TetR-expressing cell lines, TetR binds the two TetO sites and represses the CMV-MIE promoter; addition of doxycycline (Dox) releases TetR and activates GOI transcription. In cell lines lacking TetR, CMV-MIE functions as a strong constitutive promoter. **d**, Vector architecture of pHR-AIO-AB (Puro^R^ variant). A compact EF-1*α* promoter constitutively drives expression of the bicistronic open reading frame encoding rtTA-V16 and the antibiotic-resistance marker linked by a T2A peptide, providing both the transactivator and selection marker from a single transcript. The GOI is placed under the TRE3GS promoter for Dox-inducible expression. All genetic elements are flanked by unique restriction sites. **e**, Antibiotic variants of the pHR-AIO-AB system, available with puromycin, blasticidin, hygromycin or zeocin cassettes, permitting single-vector selection or orthogonal co-selection when co-infecting with multiple constructs. **f**, Inducible expression mechanism of the pHR-AIO-AB vectors. In the presence of Dox, rtTA-V16 binds TRE3GS to activate GOI transcription; in the absence of Dox, TRE3GS exhibits very low basal activity, enabling tight control of induction. **g**, Schematic overview of inducible expression after proviral integration (Puro^R^ variant). Left: The pHR-AB-CMV-TetO_2_ proviral DNA is integrated in the genome of HEK293S GnTI^−^TetR_LENTI_ or HEK293T TetR_LENTI_ cell lines constitutively expressing TetR (from a previously integrated proviral TetR–Blast^R^ cassette). TetR represses the CMV-MIE promoter until Dox is added, which releases repression and triggers GOI transcription. Constitutive expression of the EF-1*α* cassette enables antibiotic selection. Right: In regular HEK293S GnTI^−^or HEK293T cells transduced with pHR-AIO-AB vectors, the EF-1*α* cassette constitutively produces rtTA-V16 and the antibiotic-resistance marker, enabling enrichment of transduced cells and subsequent Dox-dependent induction of the GOI. Abbreviations: TKpA, thymidine kinase polyadenylation signal; CTE, constitutive transport element; Puro^R^, puromycin resistance; EF-1*α*, Human elongation factor-1 alpha promoter; cHS4, chicken hypersensitive site 4 insulator; CMV-MIE, human cytomegalovirus major immediate-early promoter; TetO_2_, two tandem Tet (tetracycline) operator sites; GOI, gene of interest; WPRE, woodchuck hepatitis virus post-transcriptional regulatory element; Zeo^R^, zeocin resistance; Hygro^R^, hygromycin resistance; Blast^R^, blasticidin resistance; TetR, Tet Repressor protein; Dox, doxycycline; T2A, *Thosea asigna* virus 2A peptide; c-myc NLS, nuclear localisation signal derived from the c-Myc protein; rtTA V16, reverse tetracycline transactivator V16; TRE3GS, minimal CMV promoter with seven TetO operator sequences; HEK293, human embryonic kidney 293; GnTI^−^, N-acetylglucosaminyltransferase I-deficient; dsDNA, double-stranded DNA.

**Table 1.**
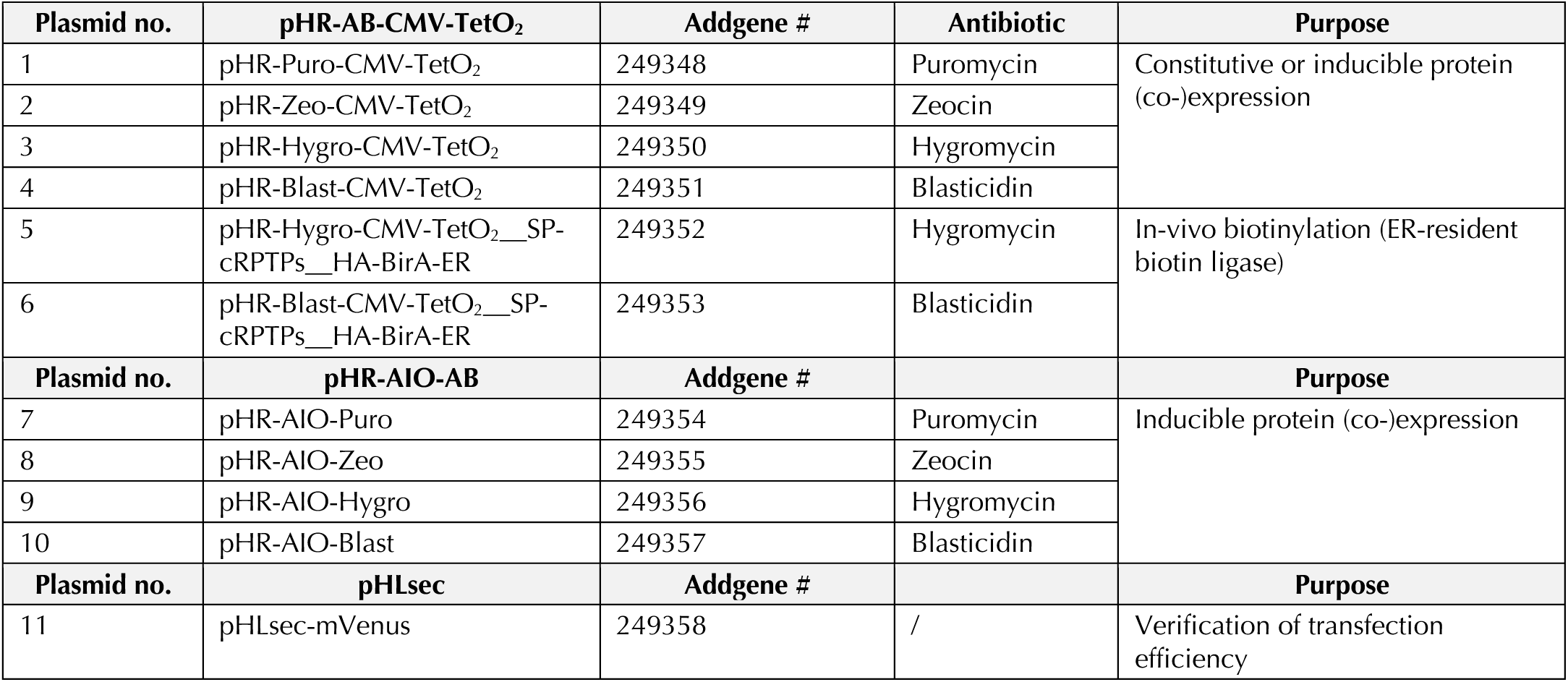
Plasmids.

**Table 2.**
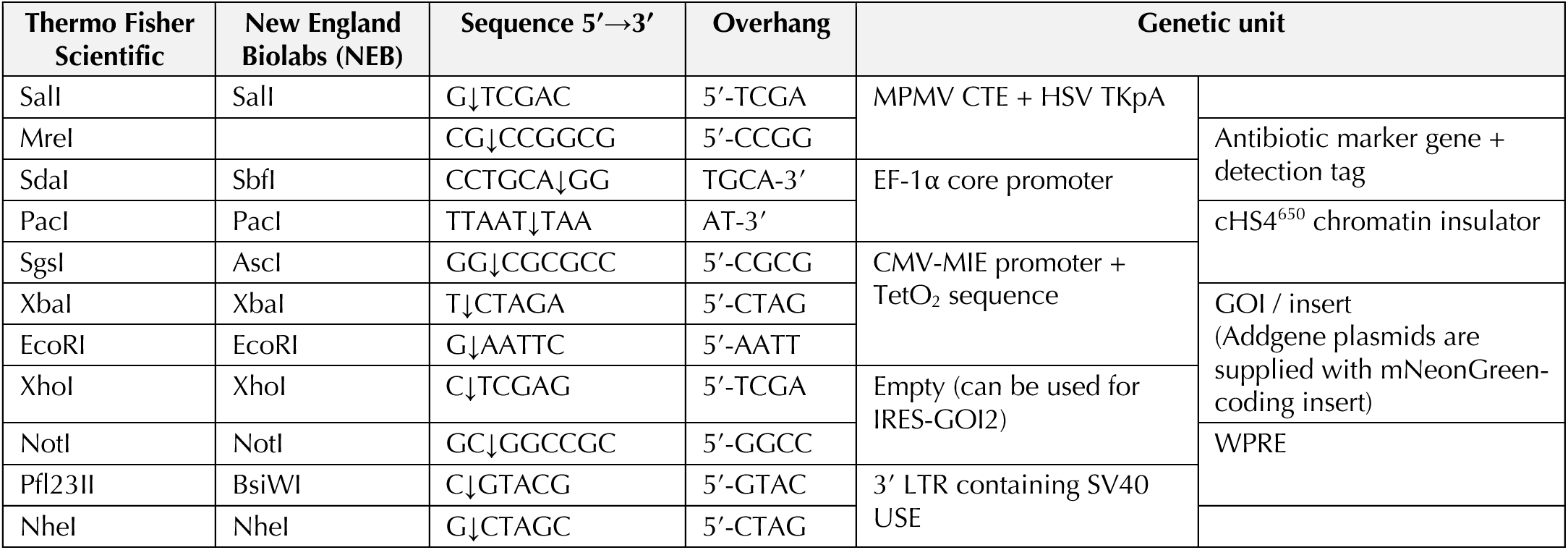
Modular arrangement and unique restriction sites in the pHR-AB-CMV-TetO_2_ transfer plasmids.

To achieve this, we added either puromycin, zeocin, hygromycin, and blasticidin selection marker cDNAs (Fig. 2b) under control of a constitutively active human elongation factor-1 alpha core (EF-1α_CORE_) promoter (*P*_EF-1_*_α_*_-CORE_). *P*_EF-1_*_α_*_-CORE_ differs from the full-length EF-1α promoter by the absence of the EF-1α intron A, and we chose it for its small size (212 bp) and ability to direct high-level protein expression in HEK293 cell lines ^2^. The transcription unit controlled by *P*_EF-1_*_α_*_-CORE_ is terminated by the Mason-Pfizer monkey virus (MPMV) RNA constitutive transport element (CTE) ^3,4^ in combination with the thymidine kinase polyadenylation signal (TKpA) (Fig. 2a).

To prevent promoter interference between *P*_CMV-MIE_ and *P*_EF-1_*_α_*_-CORE_, we converged on a design where both promoters are arranged in a “head-to-head” sense/antisense configuration and separated by the chicken hypersensitive site-4 (cHS4) chromatin insulator ^5–7^. This insulator minimises transcriptional interference between individual promoters in a dual-promoter configuration and reduces retroviral vector silencing by epigenetic modifications of integrated provirus. The 5’ 250 bp core sequence and the 3’ 400 bp sequence of the native 1200 bp cHS4 insulator recapitulate full activity ^8^ and were inserted between the *P*_CMV-TetO2_ and *P*_EF-1_*_α_*_-CORE_ as a 650 bp cHS4^650^ fragment (Fig. 2a).

We further incorporated two tandem copies of the simian vacuolating virus 40 (SV40) upstream sequence elements (USE) in the 3’ long-terminal repeat (LTR) ^9^: this leads to more stringent termination of transcription during production of the lentiviral particles. This small insertion in the 3’ LTR is unlikely to negatively impact viral titer levels ^10^. Finally, we deleted the pol/env (p/e) border fragment including the native splice acceptor (SA) site, similar to vector pLVX (Takara Bio); this modification has been shown to lead to reduced-background expression in the non-Dox-induced “off” state ^11^.

To enable inducible protein expression with the pHR-AB-CMV-TetO_2_ plasmids, we created the HEK293T TetR_LENTI_ and HEK293S GnTI^−^ TetR_LENTI_ expression cell lines (Table 3). These cell lines were themselves constructed by lentiviral transduction using the pHR-CMV_TetR-HA-NLS-P2A-BSD-Myc transfer plasmid (Addgene plasmid #113899) as previously described ^1^. They constitutively express the Tet repressor protein (TetR), which binds the TetO_2_ sites immediately downstream of the CMV-MIE promoter to block transcription (Fig. 2c). Fig. 2g schematically illustrates the mechanism of constitutive antibiotic-resistance gene expression, and TetR-mediated inducible expression.

**Table 3.**
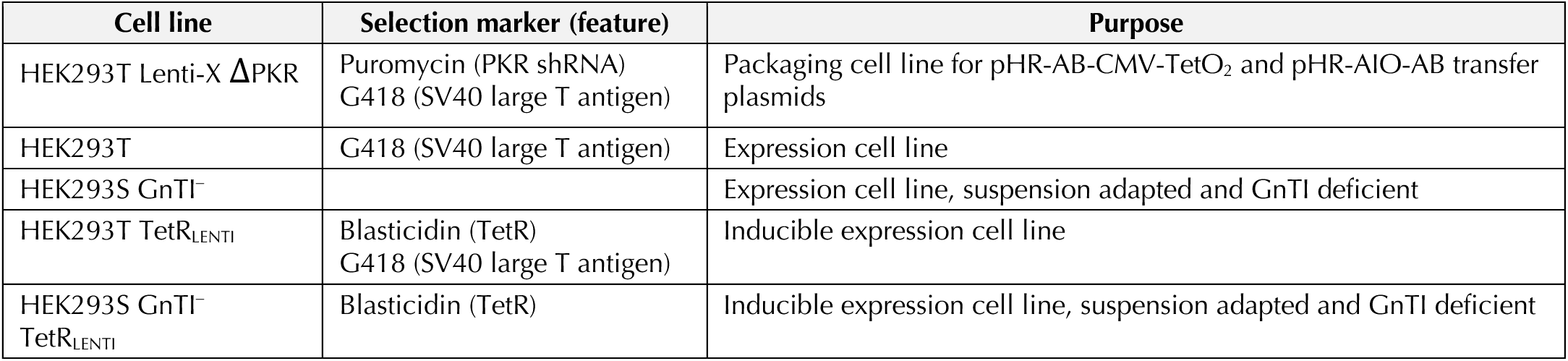
Cell lines.

### Features of the “all-in-one” pHR-AIO-AB transfer plasmids

Inducible expression using the pHR-AB-CMV-TetO_2_ system requires the use of a cell line that constitutively expresses TetR, since the *P*_CMV-MIE_ is activate in the absence of TetR (Fig. 2c). Therefore, we set out to construct a truly “all-in-one” (AIO) lentiviral expression plasmid (pHR-AIO-AB) that contains both (*i*) a constitutively expressed latest-generation reverse tetracycline transactivator (rtTA) protein together with a constitutively expressed antibiotic selection marker, as well as (*ii*) the GOI under control of the Dox-inducible TRE3GS promoter (*P*_TRE3GS_) (Fig. 2d, Table 1 and Table 4).

**Table 4.**
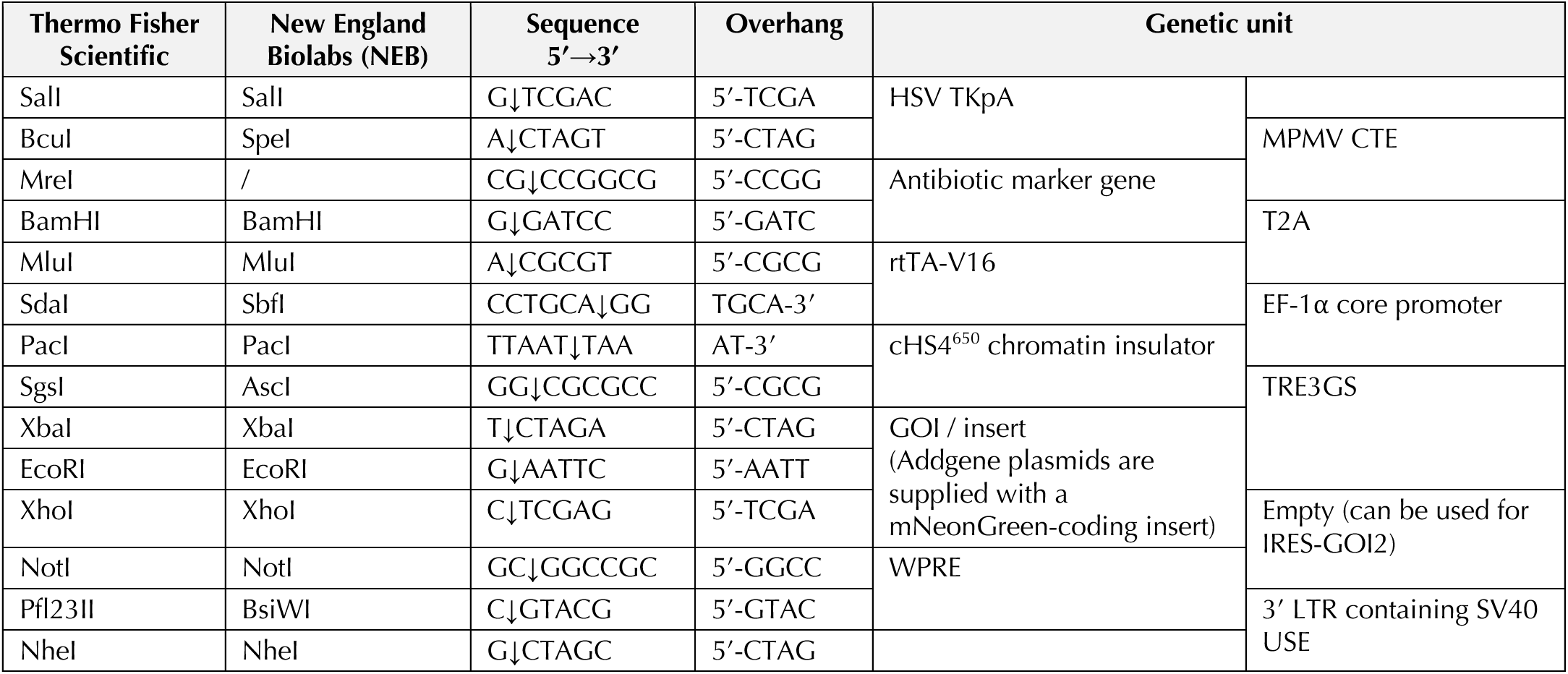
Modular arrangement and unique restriction sites in the pHR-AIO-AB transfer plasmids.

The reverse tetracycline transactivator V16 (rtTA-V16) protein is an evolved Tet-On transactivator that, in the presence of Dox, specifically binds to *P*_TRE3GS_ to activate GOI transcription and exhibits ∼100-fold higher Dox sensitivity and ∼7-fold greater activity than the original rtTA ^12,13^. It contains three minimal herpesvirus protein VP16 activation domains ^14^, and we codon-optimised the coding sequence for robust expression in HEK293 cells. To promote nuclear accumulation, we fused rtTA-V16 to a c-myc nuclear localisation signal (NLS) ^15^, and then linked it to the different antibiotic selection marker cDNAs (puromycin, zeocin, hygromycin and blasticidin) using the *Thosea asigna* virus 2A peptide (T2A) ^16^. The T2A element mediates ribosome-skipping during translation, which allows rtTA-V16 and the antibiotic-resistance protein to be produced as two separate polypeptides from a single open reading frame, and ensures equimolar expression while preserving full activity of both components. The resulting rtTA-V16-T2A-AB cassettes are placed under control of *P*_EF-1_*_α_*_-CORE_, in an arrangement identical to the pHR-AB-CMV-TetO_2_ system described above (Fig. 2e). *P*_TRE3GS_ shows very low basal expression, but high maximal expression upon Dox induction ^11,17^.

It consists of seven repeats of a 19-bp Tet operator (TetO) sequence located upstream of a minimal cytomegalovirus (CMV) promoter (Fig. 2f). *P*_TRE3GS_ is a modification of the TRE3G promoter (*P*_TRE3G_) and is optimised for higher performance in a single-vector context. *P*_TRE3GS_ is virtually silent in the absence of Dox induction since it lacks binding sites for endogenous mammalian transcription factors. Although expression from *P*_TRE3GS_ is not as strong as from *P*_CMV-MIE_, this can be advantageous for membrane proteins, where excessive expression risks saturating the secretory pathway and plasma membrane, provoking mistargeting or proteostatic stress. In practice, rtTA-V16/TRE3GS provides tight, low-leak control with tunable induction: Dox can be titrated to set expression at a level that supports proper folding and trafficking. A schematic illustrating the mechanism of constitutive antibiotic-resistance gene expression, and TetR-mediated inducible expression, is shown in Fig. 2g.

### Features of the HEK293T Lenti-X ΔPKR producer cell line

A problem with a lentiviral vector that encodes an antisense transcript (in our case, rtTA-V16 and the antibiotic marker genes) is the production of double-stranded RNAs (dsRNAs) that induce an antiviral innate immune response, such as translational repression induced by host protein kinase R (PKR; also known as Eukaryotic Translation Initiation Factor 2 Alpha Kinase 2 or EIF2AK2) ^18^. To mitigate the resulting loss of viral titer upon transfection of the transfer plasmids in the regular HEK293T Lenti-X virus production cell line ^1^, we performed short hairpin RNA (shRNA)-mediated knockdown of PKR as previously described ^19,20^ to create the HEK293T Lenti-X ΔPKR packaging cell line (Fig. 3a and Table 3). Briefly, we produced lentiviral particles encoding anti-EIF2AK2 shRNA (Sigma Aldrich MISSION shRNA TRCN0000196400) and used them to stably transduce HEK293T Lenti-X cells. Anti-EIF2AK2 shRNA expressing cells were then selected with puromycin, giving rise to the polyclonal HEK293T Lenti-X ΔPKR packaging cell line. This ΔPKR producer line consistently improves functional titre with antisense-bearing constructs and is therefore recommended for use with both the pHR-AB-CMV-TetO_2_ and pHR-AIO-AB vector suites.

**Fig. 3.**
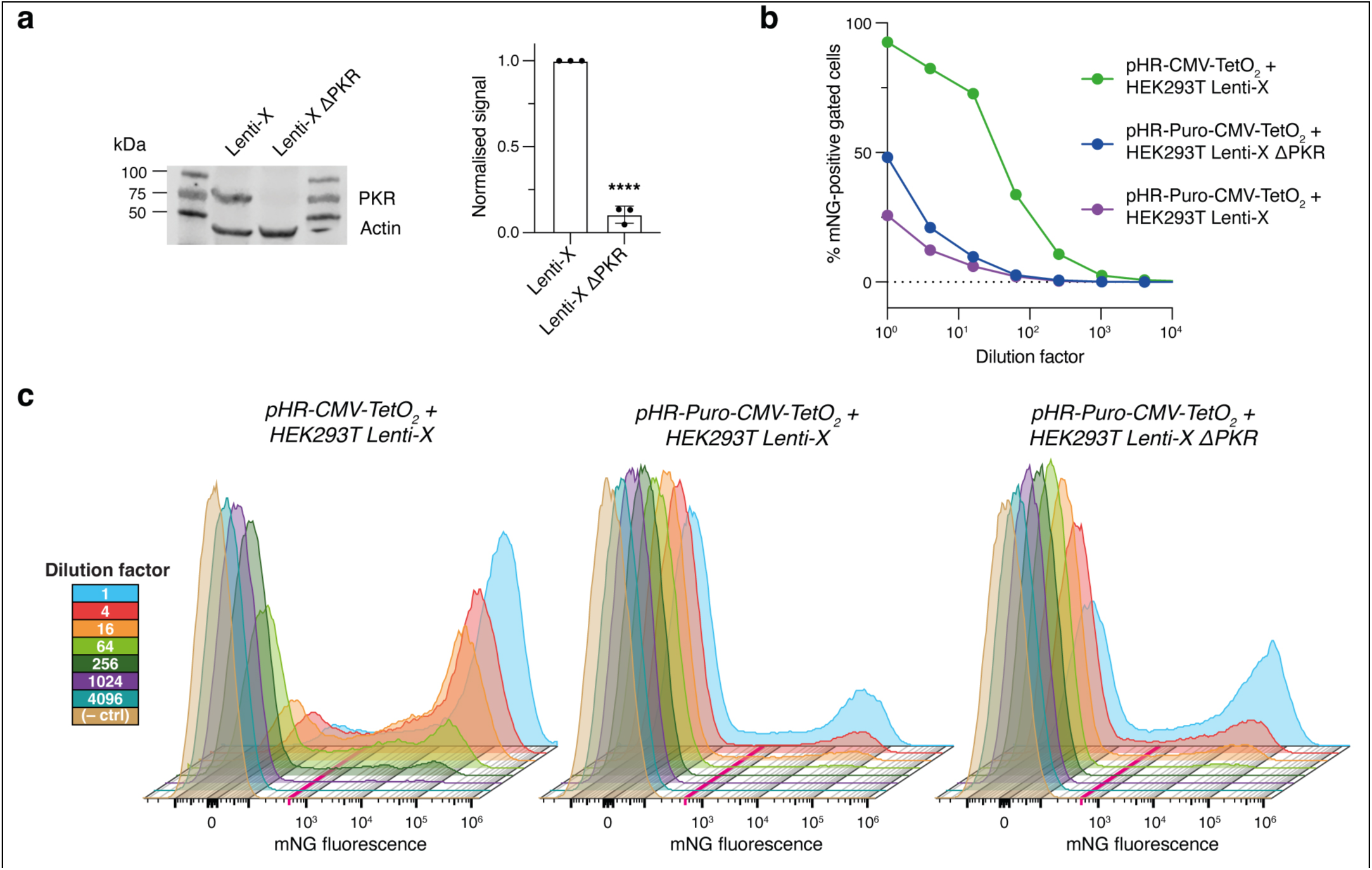
HEK293T Lenti-X ΔPKR producer cell line and functional titration of lentiviral supernatant. **a**, Western blot with quantification confirming PKR knockdown in the Lenti-X ΔPKR producer cells relative to the parental Lenti-X cell line (actin is shown as loading control). Band intensities were quantified by densitometry, normalized to actin and scaled to the Lenti-X mean (=1). Bars show mean ± s.d. of n = 3 independent experiments. Significance was tested with an unpaired two-tailed Welch’s t-test comparing Lenti-X and Lenti-X ΔPKR; **** denotes P < 0.0001. **b**, Fraction of mNeonGreen (mNG)–positive HEK293 expression cells as a function of lentiviral supernatant dilution; with the original pHR-CMV-TetO_2_ ^1^ (green), the undiluted inoculum yields ∼100% mNG-positive cells, whereas the pHR-Puro-CMV-TetO_2_ vectors (blue and purple) show lower apparent infection at the same input and therefore require antibiotic enrichment to approach uniform positivity. Virus production in Lenti-X ΔPKR cells (blue) partially mitigates this loss, increasing fraction of mNG-positive cells at a given dilution. Positivity is defined by the gate in (c). **c**, Representative flow cytometry histograms of mNG fluorescence for the dilution series shown in (b); the magenta line marks the positivity threshold used for gating (uninfected control shown for reference). Abbreviations: PKR, host protein kinase R; mNG, mNeonGreen; HEK293, human embryonic kidney 293; CMV, human cytomegalovirus; TetO₂, tandem Tet (tetracycline) operator sites; Puro, puromycin.

### Molecular cloning into the pHR-AB-CMV-TetO_2_ and pHR-AIO-AB transfer plasmids

Cloning into the pHR-AB-CMV-TetO_2_ and pHR-AIO-AB transfer plasmids is straightforward. In our hands, the simplest route is by mini-prepping the plasmid, linearising it (with EcoRI and XhoI), and assembling the PCR-amplified insert with the linearised vector using a seamless method such as In-Fusion (Takara Bio), following the manufacturer’s instructions. Although we recommend In-Fusion, the vectors are equally compatible with other routine cloning strategies including Gibson assembly, In Vivo Assembly (IVA) ^21^, and conventional restriction–ligation cloning, as long as the backbone is first linearised and the insert carries the appropriate overhangs.

Critically, do not PCR-amplify the vector backbone. These lentiviral plasmids contain repeated and highly structured elements and local regions of elevated GC content; backbone PCR therefore tends to stall polymerases and promotes slippage or recombination, yielding rearranged or truncated cassettes, poor transformation efficiency, and occasional loss of essential cis-acting sequences. For reliability and preservation of vector integrity, restrict PCR to the insert and start assembly from a clean plasmid prep of the backbone.

### Comparison to other methods

A detailed head-to-head comparison with transient transfection, monoclonal stable cell lines, piggyBac transposition and BacMam is provided in our previous protocol ^1^, and those conclusions remain unchanged with the present update. In brief, lentiviral delivery still avoids large-scale DNA preparations and cumbersome transfections, yields near-uniform expression in polyclonal populations generated in ∼3–4 weeks (versus ∼8–10 weeks for monoclonal cell lines), and offers practical advantages over BacMam for large-scale work, including stable storage of cell lines, a smaller lab footprint and no requirement for insect-cell infrastructure.

The maximum recommended lentiviral genome size is ∼8–10 kb (*i.e.*, the sequence packaged from the 5’ LTR to the end of the 3’ LTR in the transfer vector). As this limit is approached or exceeded, functional titer drops because encapsidation becomes less efficient for long RNAs. As a result, integration efficiency declines, resulting in a larger fraction of non-infectious particles. Practically, this manifests as fewer infectious units per mL and lower multiplicities of infection (MOIs), and thus requires longer selection. Multi-cistronic designs (*e.g.*, pHR-AIO-AB vectors encoding rtTA, an antibiotic-resistance gene and a GOI) therefore tend to yield lower titers than leaner constructs (*e.g.*, pHR-CMV-TetO_2_ vectors with a single expression cassette ^1^). To mitigate this limitation, our vector suites enable complex or multi-subunit assemblies to be distributed across multiple lentiviral constructs, with orthogonal antibiotic markers (puromycin, zeocin, blasticidin, hygromycin) supporting efficient co-infection and co-selection of cells that express all components.

### Biological safety

Biological safety guidance also remains exactly as described in the 2018 protocol ^1^ and is not altered by the current plasmid refinements. We continue to use a second-generation, self-inactivating (SIN) lentiviral system with transfer, envelope and packaging functions on separate plasmids, which minimises the likelihood of generating replication-competent virus; oncogenesis risk is dictated by the insert and must be assessed on a case-by-case basis. Operational risk-reduction practices are unchanged (no routine concentration of viral supernatant; all work in a Class II MSC with appropriate personal protective equipment, decontamination and waste handling), and experiments are performed under biosafety level 2 BSL-2 conditions.

## Materials

### Biological materials

**! CAUTION** The cell lines used in your research should be regularly checked to ensure they are authentic and are not infected with mycoplasma.

- HEK293T Lenti-X ΔPKR cells (G418^R^, Puro^R^; obtainable by request to J.E.)
- HEK293T cells (G418^R^; ATCC, cat. no. CRL-3216)
- HEK293S GnTI^−^ cells (ATCC, cat. no. CRL-3022)
- HEK293T TetR_LENTI_ cells (G418^R^, Blast^R^; obtainable by request to J.E.)
- HEK293S GnTI^−^ TetR_LENTI_ cells (Blast^R^; obtainable by request to J.E.)

### Reagents

- Gibco FBS (Thermo Fisher Scientific, cat. no. 10270106)
- Gibco MEM non-essential amino acids solution (NEAA; Thermo Fisher Scientific, cat. no. 11140-035)
- Gibco PBS (calcium- and magnesium-free; Thermo Fisher Scientific, cat. no. 10010015)
- Gibco trypsin-EDTA (0.05% (wt/vol), phenol red; Thermo Fisher Scientific, cat. no. 25300054)
- Gibco DMEM/Nutrient Mixture F-12 (DMEM/F-12; with added L-Gln; Thermo Fisher Scientific, cat. no. 11320-074)
- Polyethylenimine (PEI; 25 kDa, branched; Sigma Aldrich, cat. no. 408727)
- Polybrene infection reagent (10 mg/mL stock = 1,000×; Sigma Aldrich, cat. no. TR-1003-G)
- Doxycycline hydrochloride (Dox; Sigma Aldrich, cat. no. D3447)
- Blasticidin S hydrochloride (10 mg/mL; Thermo Fisher Scientific, cat. no. A1113903)
- Zeocin (100 mg/mL; Thermo Fisher Scientific, cat. no. R25001)
- Hygromycin B (Sigma Aldrich cat. no. H3274)
- Puromycin dihydrochloride (10 mg/mL; Thermo Fisher Scientific, cat. no. A1113802)
- Dimethyl sulfoxide (DMSO; sterile-filtered, suitable for cell culture; Sigma Aldrich, cat. no. D2650)
- Rely+On Virkon disinfectant powder (VWR, cat. no. 148-0202)
- Microsol 4 decontaminant (Anachem, cat. no. 30312915)
- (D-)biotin (Merck, cat. no. B4639)

### Sequencing primers

- pHR-AB-CMV-TetO_2_ forward sequencing primer AB-FWD (5ʹ-CGTCGTCGTGCTCGTTTAGTG-3ʹ)
- pHR-AIO-AB forward sequencing primer AIO-FWD (5ʹ-GAGTAAACTTCAATTCCACAACAC-3ʹ)
- pHR-AB-CMV-TetO_2_, pHR-AIO-AB reverse sequencing primer AB-REV, AIO-REV (5ʹ-GAGCAACATAGTTAAGAATACCAGTC-3ʹ)

### Plasmids

**▴CRITICAL** All transfer plasmids are available through the non-profit Addgene plasmid repository and are subject to a Uniform Biological Material Transfer Agreement (UBMTA).

- Transfer plasmid: pHR-AB-CMV-TetO_2_/pHR-AIO-AB plasmids (Addgene catalogue numbers are listed in Table 1)
- pMD2.G lentiviral envelope plasmid (Addgene, cat. no. 12259)
- psPAX2 lentiviral packaging plasmid (Addgene, cat. no. 12260)

### Equipment

- Class II microbiological safety cabinet (MSC; Thermo Scientific, model no. Herasafe KS 12)
- Long-cuff nitrile gloves (StarGuard, PROTECT+ model)
- CO_2_ incubator (Thermo Scientific, model no. HERAcell 150i)
- Tissue culture plate (6-well plates, flat bottom; Greiner Bio-One, cat. no. 657160)
- Tissue culture plate (T75, filter cap, 75 cm^2^; Greiner Bio-One, cat. no. 658175)
- Tissue culture plate (T175, filter cap, 175 cm^2^; Greiner Bio-One, cat. no. 660175)
- Sterile 30-mL Luer-lock plastic syringes BD Plastipak (Becton Dickinson, cat no. 301229)
- Syringe filter unit, polyethersulfone (PES; sterile, 0.45 μm; Sigma Aldrich, cat. no. SLHPM33RS)
- Syringe filter unit, polyethersulfone (PES; sterile, 0.22 μm; Sigma Aldrich, cat. no. SE2M228I04)
- Polystyrene disposable standard serological pipettes (5 mL; Greiner Bio-One, cat. no. 606180)
- Polystyrene disposable standard serological pipettes (10 mL; Greiner Bio-One, cat. no. 607180)
- Polystyrene disposable standard serological pipettes (25 mL; Greiner Bio-One, cat. no.760180)
- Polystyrene disposable standard serological pipettes (50 mL; Greiner Bio-One, cat. no. 768180)
- Bottle-top filter units (Steritop, 500 mL, sterile, 45-mm threaded; Sigma Aldrich, cat no. SCGPT05RE)
- CryoPure cell culture cryogenic tubes (1.8 mL; Sarstedt cat. no. 72.379.002)
- Controlled-rate freezing container (CoolCell LX; BioCision, cat. no. BCS-405G)

### Reagent setup

#### DMEM/F-12/10% FBS

Add 50 mL of FBS (to obtain 10% (vol/vol)) and 5 mL of NEAA (to obtain 1% (vol/vol)) to 445 mL of DMEM/F-12. Store the medium at 4 °C and, immediately before use, visually inspect it (clarity, colour, precipitate) to confirm there are no signs of contamination.

#### DMEM/F-12/2% FBS

Add 10 mL of FBS (to obtain 2% (vol/vol)) and 5 mL of NEAA (to obtain 1% (vol/vol)) to 485 mL of DMEM/F-12. Store the medium at 4 °C and, immediately before use, visually inspect it (clarity, colour, precipitate) to confirm there are no signs of contamination.

#### DMEM/F-12/SFM (serum-free medium)

Add 5 mL of NEAA (to obtain 1% (vol/vol)) to 495 mL of DMEM/F-12. Store the medium at 4 °C and, immediately before use, visually inspect it (clarity, colour, precipitate) to confirm there are no signs of contamination.

#### PEI (1 mg/mL stock)

The PEI stock solution is highly viscous and cannot be pipetted. First, pour and weigh 1–5 mL straight from the bottle into a 50-mL tube. PEI has a density of 1.030 g/mL at 25 °C. Add ultrapure water to prepare a 100 mg/mL stock solution. Rotate the tube overnight to mix. Once the solution is homogeneous, dilute it to 1 mg/mL in ultrapure water and adjust the pH to 7.0 with hydrochloric acid (HCl). Filter-sterilise, using a 0.2-μm syringe filter unit inside a biological safety cabinet. Make aliquots and store at −20 °C for up to 1 year.

#### Dox (10 mg/mL stock)

Dissolve 50 mg in 100% (vol/vol) ethanol to a final volume of 5 mL and filter-sterilize, using a 0.2-μm syringe filter unit inside a biological safety cabinet. Make aliquots and store at −20 °C for up to 8 weeks.

**▴CRITICAL** Do not expose Dox to direct sunlight.

#### Hygromycin (50 mg/mL stock)

Dissolve 500 mg in ultrapure water to a final volume of 10 mL and filter-sterilise, using a 0.2-μm syringe filter unit. Make aliquots and store at –20 °C for up to 8 weeks.

**! CAUTION** Hygromycin is toxic. Always wear PPE: laboratory coat, safety eyewear and disposable long-cuff gloves. Weigh the powder in a chemical safety cabinet and prepare the solution inside a biological safety cabinet.

#### Purification of plasmid DNA

Purify plasmid DNA, using a QIAprep Spin Miniprep Kit (Qiagen) or equivalent kit from a different brand, according to the manufacturer’s instructions. Plasmid can be stored indefinitely at –20 °C.

#### Growth and maintenance of adherent and suspension HEK293-derived cells

Culture cells as described in detail in our previous protocol ^1^.

**! CAUTION** Cell cultures are a potential biological hazard. Working with HEK293 cells requires BSL2 practices. Perform the work in an approved laminar flow cabinet, using sterile techniques, and comply with the appropriate institutional biosafety guidelines. This includes wearing protective clothing and eyewear, cleaning of working surfaces, and proper disposal of waste before and after performing experiments.

## Procedure

### Seeding of HEK293T Lenti-X ΔPKR producer cells (Day 1) ● Timing ∼3.5 d

1. In the afternoon, add ∼3.1 × 10^6^ HEK293T trypsinised Lenti-X ΔPKR cells (equivalent to one-sixth of the total cells recovered after trypsinising a confluent T75 flask ^1^) in 20 mL of DMEM/F-12/10% FBS medium to a fresh T75 flask, bringing it to ∼16% confluency. Cell culture is described in detail in our previous protocol ^1^.

2. Incubate the flask for 3.5 d at 37 °C in a humidified incubator with 5% CO_2_.

**▴CRITICAL STEP** Lentivirus can be produced from regular HEK293T and HEK293 Lenti-X cells, but the HEK293 Lenti-X ΔPKR cell line has been engineered to yield viral titers higher than those of HEK293T cells in combination with the pHR-AB-CMV-TetO_2_ and pHR-AIO-AB transfer plasmids (Fig. 3b,c).

**▲CRITICAL STEP** The health of the HEK293T Lenti-X ΔPKR producer cell line and of the HEK293 expression cell lines is critical for obtaining good lentiviral titers, efficient transduction and robust protein expression. Maintain them diligently and split the cells 1/6 twice a week at fixed times; we suggest Monday morning and Thursday afternoon. We do not use the cells beyond passage 20 (P20; ∼10 weeks of culture) to ensure maximum viability and viral yield. When starting the HEK293T Lenti-X ΔPKR cell line from a frozen stock, we advise to passage them at least once to ensure proper recovery before seeding them for transfection.

**! CAUTION** Lentiviral particles are a potential biological hazard and require biosafety level 2 (BSL2) or 2+ (BSL2+) practices, depending on the institutional and governmental biosafety guidelines. Perform the work in an approved laminar flow cabinet using sterile techniques and comply with the appropriate institutional biosafety guidelines. A list of risk reduction measures can be found in the “Biological Safety Considerations” section in our previous protocol ^1^.

**? TROUBLESHOOTING**

### Seeding of HEK293 expression cells (Day 1) ● Timing ∼3.5 d

3. In the afternoon, add ∼1.9 x 10^6^ (HEK293T; equivalent to one-tenth of the total cells recovered after trypsinising a confluent T75 flask) or ∼3.1 × 10^6^ (HEK293S; equivalent to one-sixth of the total cells recovered after trypsinising a confluent T75 flask) expression cells in 20 mL of DMEM/F-12/10% FBS medium to a new T75 flask.

| Expression cell line | Transfer plasmid used for transduction | Specifics of the transduced expression cell line |
| --- | --- | --- |
| HEK293T | pHR-AIO-AB | Inducible expression, adherent format |
|  | pHR-AB-CMV-TetO<sub>2</sub> | Constitutive expression, adherent format |
| HEK293S GnT1<sup>-</sup> | pHR-AIO-AB | Inducible expression, adherent or suspension format |
|  | pHR-AB-CMV-TetO<sub>2</sub> | Constitutive expression, adherent or suspension format |
| HEK293T TetR<sub>LENTI</sub> | pHR-AB-CMV-TetO<sub>2</sub> | Inducible expression, adherent format |
| HEK293S GnT1<sup>-</sup> TetR<sub>LENTI</sub> | pHR-AB-CMV-TetO<sub>2</sub> | Inducible expression, adherent or suspension format |

4. (Optional) In the case of HEK293T TetR_LENTI_ or HEK293S GnTI^−^ TetR_LENTI_ cells, add blasticidin (final concentration of 10 μg/mL from a 10 mg/mL 1,000× stock solution) to the DMEM/F-12/10% FBS medium, to maintain selective pressure and ensure continued TetR expression.

5. Incubate the flask for 3.5 d at 37 °C in a humidified incubator with 5% CO_2_.

### Transfection of HEK293T Lenti-X ΔPKR cells (Day 5) ● Timing ∼2.5 d

6. In the morning, prepare the following plasmid DNA transfection mix (33 μg of total DNA; 1:1:1:0.3 (wt/wt/wt/wt) transfer/packaging/envelope/mVenus plasmid ratio) in a sterile 1.5-mL tube:

- 10 μg pHR-AB-CMV-TetO_2_ or pHR-AIO-AB transfer plasmid variant encoding the GOI
- 10 μg psPAX2 packaging plasmid
- 10 μg pMD2.G envelope plasmid
- 3 μg pHLsec-mVenus transfection control plasmid

(mVenus is included as a fluorescent marker of producer-cell transfection efficiency, which can quickly be assessed on a standard wide-field fluorescence microscope.)

Top up the transfection mix with DMEM/F-12/SFM medium to a total volume of 0.25 mL and gently mix the suspension.

**? TROUBLESHOOTING**

7. Prepare 82.5 μL of PEI (1:2.5 (wt/wt) DNA:PEI ratio) in a sterile 1.5-mL tube. Top up with DMEM/F-12/SFM medium to a total volume of 0.25 mL, and gently mix the suspension.

8. When both solutions are ready, add the 0.25 mL of PEI mix to the 0.25 mL of plasmid DNA transfection mix for a total of 0.5 mL in a 1.5-mL tube, and gently mix the suspension.

**▲CRITICAL STEP** In our experience, both the volume in which the DNA/PEI transfection mix is prepared, and the DNA/PEI ratio, have a great effect on the transfection efficiency. We recommend adhering to the values stated in Steps 7–8.

9. Centrifuge the 1.5-mL tube briefly at low speed (100 g, 22–24 °C, 30 s) to collect the liquid at the bottom of the tube, and then leave it for incubation in the flow cabinet for 15–20 min.

10. The T75 flask with HEK293T Lenti-X cells should now be ∼70–90% confluent (continued from Step 1); take off and discard the growth medium; wash with 10 mL PBS and replace with 19.5 mL DMEM/F-12/2%FBS medium.

11. Add the 0.5 mL DNA/PEI mix to the 19.5 mL expression medium for a total of 20 mL and gently tilt the flask to distribute the DNA/PEI mix and cover all cells.

12. Incubate the T75 flask for 2.5 d at 37 °C in a humidified incubator with 5% CO_2_ to allow for transfection and the ensuing production of viral particles.

### Passaging of expression cells (Day 5) ● Timing ∼2.5 d

13. In the morning, add ∼3.1 x 10^6^ (∼1/6 split; HEK293T) or ∼4.7 x 10^6^ (∼1/4 split; HEK293S) trypsinised expression cells in 20 mL of DMEM/F-12/10% FBS medium to a new T75 flask.

14. (Optional) In the case of HEK293T TetR_LENTI_ or HEK293S GnTI^−^ TetR_LENTI_ cells, add blasticidin (final concentration of 10 μg/mL from a 10 mg/mL 1,000× stock solution) to the DMEM/F-12/10% FBS medium.

15. Incubate the flask for 2.5 d at 37 °C in a humidified incubator with 5% CO_2_.

### Collection of the lentivirus-containing medium (Day 7) ● Timing ∼30 min

16. In the next steps (Steps 17–20), a single 30 mL inoculum is prepared to infect the expression cells. For single-virus infections, 20 mL of filtered virus-containing medium is topped up with 10 mL fresh medium. For co-infection with multiple viral preparations (for example GOI1–puromycin, GOI2–zeocin and GOI3–hygromycin), separately-filtered virus-containing media are simply combined: 2 × 10 mL for two viruses or 3 × 6.5 mL for three, and then topped up with 10 mL fresh medium.

|  | GOI 1 | GOI 2 | GOI 3 | Fresh DMEM medium |
| --- | --- | --- | --- | --- |
| Single-virus infection | 20 mL | / | / | 10 mL |
| Double-virus infections | 10 mL | 10 mL | / | 10 mL |
| Triple-virus infections | 6.5 mL | 6.5 mL | 6.5 mL | 10 mL |

17. In the afternoon, collect the required amount (as specified above) of conditioned medium from (each of) the HEK293T Lenti-X ΔPKR T75 flask(s) into a (or multiple) sterile 50-mL tube(s).

18. Filter each conditioned medium through a 0.45-μm filter unit attached to a Luer-lock syringe into a new sterile 50-mL tube to remove cell debris.

**! CAUTION** Handle the medium containing lentiviral particles with great care to avoid spills. Use a stable rack to support the 50 mL tube, ensure that the filter unit is properly screwed onto the syringe, and keep one hand supporting the syringe until all medium is pushed safely through the filter.

**▴CRITICAL STEP** Do not use a 0.22-μm filter unit, because this may shear the viral particles. Use only cellulose acetate or polyethersulfone (PES) low-protein-binding filters. Avoid nitrocellulose filters, which bind lentiviral envelope proteins and destroy the virus.

19. For both single-virus or multiple-virus infections, add 10 mL of fresh DMEM/F-12/10% FBS medium to the 50-mL tube, yielding a total of 30 mL.

20. Add 30 μL of polybrene (hexadimethrine bromide; from a 10 mg/mL, 1,000× stock solution) to the 30 mL lentivirus-containing medium and mix gently. Polybrene is a cationic polymer that reduces charge repulsion between viral particles and the cell membrane ^22^, thereby enhancing transduction efficiencies by promoting virus–host cell fusion.

**? TROUBLESHOOTING**

### Infection of the expression cells (Day 7) ● Timing ∼2 d

21. The T75 flask with expression cells (continued from Step 15) is now ∼80–90% confluent; remove and discard the DMEM/F-12/10% FBS medium; wash with 10 mL of PBS and replace the medium with the 30 mL of filtered lentivirus-containing medium (continued from Step 20).

22. Incubate the T75 flask at 37 °C in a humidified incubator with 5% CO_2_ for 2 d. The viral particles will now infect the cells and stably integrate their genetic material into the host cell genome, generating a polyclonal stable cell line.

**? TROUBLESHOOTING**

### Passaging of the transduced expression cells (Day 9) ● Timing ∼2.5 d

23. The T75 flask will now be ∼100% confluent. In the afternoon, remove and discard the old medium, wash the transduced cells 3 times with PBS, and subsequently split the cells to two new T75 flasks: a 1/10 and 1/15 split (for HEK293T cells), or a 1/8 and 1/10 split (for HEK293S cells).

24. Incubate the T75 flasks for 2.5 d at 37 °C in a humidified incubator with 5% CO_2_. One of the flasks will be used to start the antibiotic selection procedure and the other one as a backup (and to cryopreserve a “before-selection” sample of the transduced cells).

**▲CRITICAL STEP** The transduced expression cells are passaged before starting the antibiotic selection to allow for recovery and for expression of the antibiotic resistance gene(s), as wells as to obtain a sub-confluent (and actively dividing) population on which the subsequent antibiotic treatment should be most effective. As the growth and recovery of the cells after transduction tends to be variable, we prefer splitting at a sufficiently high dilution to ensure obtaining a sub-confluent monolayer (*i.e.* a single, contiguous layer of cells at ∼50% confluency) after 2.5 d incubation.

### Start of polyclonal stable cell line selection by antibiotic treatment of the transduced expression cells (Day 12) ● Timing ∼2 d

25. In the morning, use the T75 flask that should now be ∼50–60% confluent to start the antibiotic selection. Replace the medium on the cells with 20 mL fresh DMEM/F-12/10% FBS medium containing the desired antibiotic concentration.

| Antibiotic stock concentration | Puromycin | Blasticidin | Hygromycin | Zeocin |
| --- | --- | --- | --- | --- |
| Stock concentration | 10 mg/mL | 10 mg/mL | 50 mg/mL | 100 mg/mL |
| Final concentration | 10 µg/mL | 10 µg/mL | 200–400 µg/mL | 200–400 µg/mL |
| µL of stock per 20 mL medium | 20 µL | 20 µL | 80–160 µL | 40–80 µL |

~~~
26. Incubate the T75 flask for 2 d at 37 °C in a humidified incubator with 5% CO_2_.
~~~

**! CAUTION** The antibiotics (puromycin, blasticidin, hygromycin and zeocin) used in the selection procedure are toxic. Always wear PPE: laboratory coat, safety eyewear and disposable gloves.

▲**CRITICAL STEP** These working concentrations are the levels we routinely use to select HEK293-derived lines, based on repeated kill curves and the robust expression of the antibiotic-resistance cassettes from our lentiviral transfer vectors. At the indicated final concentrations (10 µg/mL puromycin, 10 µg/mL blasticidin, 200–400 µg/mL hygromycin, and 200–400 µg/mL zeocin), non-transduced cells typically round up, bleb, and detach within 24–48 h, while transduced populations remain viable under routine medium changes.

▴**CRITICAL STEP** The optimal dose, antibiotic exposure time, and number of selection cycles may vary with the antibiotic, the expression cell line (and its health) and with MOI. Monitor cultures daily and adjust the selection course to the observed response. We typically incubate the cells 1–2 d with antibiotic and observe <50% cell death, indicative of a robust viral titre and successful infection. If >50% cell death should be observed, remove the media containing dead cells and replace it with fresh DMEM/F-12/10% FBS medium without antibiotics for 1–2 d, to allow recovery. The surface space freed up by detachment of the dead cells will allow the surviving cells to expand.

▴**CRITICAL STEP** In our hands, puromycin acts fastest, followed by blasticidin and hygromycin, with zeocin typically the slowest. As a result, for single-vector infections we generally prefer puromycin to achieve rapid yet robust selection.

▲**CRITICAL STEP** When co-infecting with two or more lentiviral vectors, greater cell loss is expected during multi-antibiotic selection, as only cells that receive all required vectors will survive. Under a Poisson model, the likelihood that a cell receives at least one copy of a given viral vector is P(≥1) = 1 – e^−MOI^. For co-infection with multiple independent vectors, the probability that a cell receives all required constructs is the product of these individual probabilities, assuming independent infection events. As shown in the table, at lower MOI values this probability declines sharply with each additional vector. For example, assuming a MOI of 0.5 for each GOI in the final inoculum, when cells are transduced with one GOI, ∼39% are expected to be infected; with two GOIs, ∼15% of cells will receive both vectors; with three GOIs ∼6% will receive all three; and with four GOIs, only ∼2% will receive all four. To improve yield without excessively increasing copy number, use modestly higher MOIs (e.g., 1–2 per vector), consider sequential infections with selection between rounds, maintain a backup culture, and adjust antibiotic exposure based on daily monitoring.

|  | MOI = 0.5 | MOI = 1 | MOI = 2 | MOI = 3 |
| --- | --- | --- | --- | --- |
| One GOI | 39% | 63% | 86% | 95% |
| Two GOIs | 15% | 40% | 75% | 90% |
| Three GOIs | 6% | 25% | 65% | 86% |
| Four GOIs | 2% | 16% | 56% | 82% |

**? TROUBLESHOOTING**

### Passaging of “backup” cells and cryopreservation of non-selected transduced cells (Day 12) ● Timing ∼3.5 d

27. In the morning, if the confluency of the cells in the “backup” T75 flask is >80%, these cells can be split 1/10 (HEK293T) or 1/6 (HEK293S) to a new T75 flask, and the remaining trypsinised cell suspension can be used for cryopreservation (as described in our original protocol ^1^).

28. Incubate the new “backup” T75 flask at 37 °C in a humidified incubator with 5% CO_2_ for 3.5 d. Maintain this backup flask as long as required by splitting it every 3.5 d.

**▲CRITICAL STEP** Maintain a parallel backup flask of transduced cells without antibiotics until it is confirmed that the selection regime behaves as expected (predictable kill kinetics and healthy regrowth). This allows a rapid restart if the selected culture declines, becomes contaminated, or needs re-optimisation (antibiotic dose/exposure cycles). In addition, cryopreservation of aliquots of the non-selected population acts as a longer-term safeguard.

### Passaging of antibiotic-selected cells (Day 14) ● Timing ∼2.5 d

29. In the morning, after 2 d of incubation with antibiotic, dead cells should have detached from the flask surface and be floating in the medium. Wash the cells twice gently with 10 mL of PBS to remove the dead cells, and split the cells 1/10 (HEK293T) or 1/6 (HEK293S) into a new T75 flask, using fresh DMEM/F-12/10% FBS medium without antibiotics.

30. Incubate the T75 flask at 37 °C in a humidified incubator with 5% CO_2_. Grow the cells (typically for 2.5 d) until they reach 50–60% confluency, at which point a new round of antibiotic treatment can be started on the adherent cells.

**▲CRITICAL STEP** The selected expression cells are passaged before starting the second antibiotic treatment to allow for recovery and to ensure that selection begins on a sub-confluent, actively dividing population. Adjust the incubation time as needed to obtain a sub-confluent monolayer (*i.e.* a single, contiguous layer of cells at ∼50–60% confluency).

**? TROUBLESHOOTING**

### Second round of antibiotic selection (Day 16) ● Timing ∼2.5 d

31. In the afternoon, the T75 flask should be ∼50–60% confluent and ready to start the antibiotic selection. Replace the medium on the cells with 20 mL fresh DMEM/F-12/10% FBS medium containing the desired antibiotic concentration, as in Step 25.

32. Incubate the T75 flask for 2.5 d at 37 °C in a humidified incubator with 5% CO_2_.

### Passaging of antibiotic-selected cells (Day 19) ● Timing ∼3.5 d

33. In the morning, after 2.5 d of incubation with antibiotic, any dead cells should have again detached from the flask surface and be floating in the medium. Wash the cells twice gently with 10 mL of PBS to remove the dead cells, and split the cells 1/10 (HEK293T) or 1/6 (HEK293S) into a new T75 flask, using fresh DMEM/F-12/10% FBS medium without antibiotics.

34. Incubate the T75 flask at 37 °C in a humidified incubator with 5% CO_2_. Grow the cells (typically for 3–3.5 d) until they reach a confluency of 80–100%, at which point they can be used for cryopreservation and/or expanded for small-scale expression tests or large-scale protein production.

▲**CRITICAL STEP** In our experience, we see very little additional cell death after the second round of antibiotic selection, and hence we consider the population to be fully selected and enriched. If not, continue with one or more additional cycles of splitting and antibiotic selection until the culture regrows robustly and no more cell death is observed.

◼**PAUSE POINT** The enriched cell line can be trypsinised and long-term stored in LN_2_ for future use, as described in detail in our previous protocol ^1^.

### Expansion and induction of selected cells for (large-scale) protein production (Day 22) ● Timing ∼2–3 weeks

35. At this stage, the selected, inducible population can be expanded in the preferred format: adherent (*e.g.*, T-flasks, cell factories, HYPERFlasks, roller bottles) or suspension (shake flasks, wave bags, bioreactors), and protein expression can be induced at the desired time point with Dox. Induction will depend on the chosen vector/cell-line pairing; pHR-AB-CMV-TetO_2_ enables inducible expression in TetR-expressing lines or constitutive expression in regular HEK293 cells, whereas pHR-AIO-AB provides a true single-vector inducible system. For detailed guidance on scale-up, media and exchange regimes, Dox induction parameters, timing, and harvest/processing for soluble versus membrane targets, we refer to our previous protocol ^1^.

▴**CRITICAL STEP** For soluble proteins, we typically perform expression at 37 °C for 5 d using 1 µg/mL Dox (for the HEK293 target strains described here), but the optimal parameters for maximum yield and correct folding should be determined empirically for each target. The newly designed pHR-AIO-AB system provides tighter control over expression levels (Fig. 5b), which is advantageous for proteins that are difficult to fold or burdensome/toxic to cells.

### Troubleshooting

Troubleshooting advice can be found in Table 5.

**Table 5.**
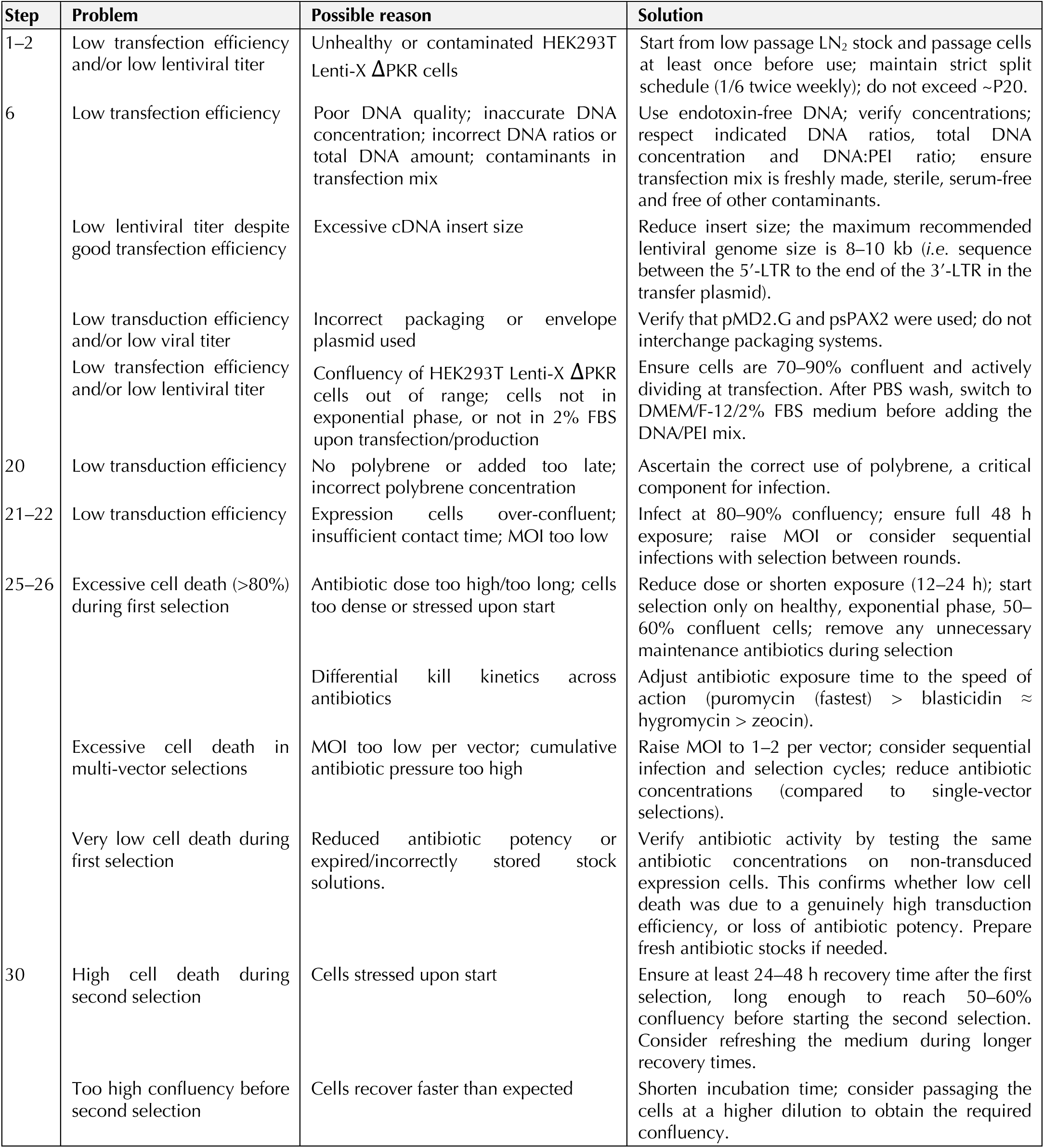
Troubleshooting table.

| Step | Problem | Possible reason | Solution |
| --- | --- | --- | --- |
| 1–2 | Low transfection efficiency and/or low lentiviral titer | Unhealthy or contaminated HEK293T Lenti-X ΔPKR cells | Start from low passage LN<sub>2</sub> stock and passage cells at least once before use; maintain strict split schedule (1/6 twice weekly); do not exceed ~P20. |
| 6 | Low transfection efficiency | Poor DNA quality; inaccurate DNA concentration; incorrect DNA ratios or total DNA amount; contaminants in transfection mix | Use endotoxin-free DNA; verify concentrations; respect indicated DNA ratios, total DNA concentration and DNA:PEI ratio; ensure transfection mix is freshly made, sterile, serum-free and free of other contaminants. |
|  | Low lentiviral titer despite good transfection efficiency | Excessive cDNA insert size | Reduce insert size; the maximum recommended lentiviral genome size is 8–10 kb (*i.e.* sequence between the 5'-LTR to the end of the 3'-LTR in the transfer plasmid). |
|  | Low transduction efficiency and/or low viral titer | Incorrect packaging or envelope plasmid used | Verify that pMD2.G and psPAX2 were used; do not interchange packaging systems. |
| | Low transfection efficiency and/or low lentiviral titer | Confluency of HEK293T Lenti-X ΔPKR cells out of range; cells not in exponential phase, or not in 2% FBS upon transfection/production | Ensure cells are 70–90% confluent and actively dividing at transfection. After PBS wash, switch to DMEM/F-12/2% FBS medium before adding the DNA/PEI mix. |
| 20 | Low transduction efficiency | No polybrene or added too late; incorrect polybrene concentration | Ascertain the correct use of polybrene, a critical component for infection. |
| 21–22 | Low transduction efficiency | Expression cells over-confluent; insufficient contact time; MOI too low | Infect at 80–90% confluency; ensure full 48 h exposure; raise MOI or consider sequential infections with selection between rounds. |
| 25–26 | Excessive cell death (>80%) during first selection | Antibiotic dose too high/too long; cells too dense or stressed upon start | Reduce dose or shorten exposure (12–24 h); start selection only on healthy, exponential phase, 50–60% confluent cells; remove any unnecessary maintenance antibiotics during selection |
| | | Differential kill kinetics across antibiotics | Adjust antibiotic exposure time to the speed of action (puromycin (fastest) > blasticidin ≈ hygromycin > zeocin). |
|  | Excessive cell death in multi-vector selections | MOI too low per vector; cumulative antibiotic pressure too high | Raise MOI to 1–2 per vector; consider sequential infection and selection cycles; reduce antibiotic concentrations (compared to single-vector selections). |
|  | Very low cell death during first selection | Reduced antibiotic potency or expired/incorrectly stored stock solutions. | Verify antibiotic activity by testing the same antibiotic concentrations on non-transduced expression cells. This confirms whether low cell death was due to a genuinely high transduction efficiency, or loss of antibiotic potency. Prepare fresh antibiotic stocks if needed. |
| 30 | High cell death during second selection | Cells stressed upon start | Ensure at least 24–48 h recovery time after the first selection, long enough to reach 50–60% confluency before starting the second selection. Consider refreshing the medium during longer recovery times. |
|  | Too high confluency before second selection | Cells recover faster than expected | Shorten incubation time; consider passaging the cells at a higher dilution to obtain the required confluency. |

### Timing

The entire protocol, starting from the transfection of Lenti-X cells to obtaining a fully antibiotic-selected transduced expression cell line ready for expansion, takes ∼20 d. The timing for each stage of the Procedure is summarised below.

Step 1–2 and 3–5 (Day 1), HEK293 producer and expression cell seeding: ∼3.5 d

Steps 6–12 (Day 5), HEK293T Lenti-X cell transfection: ∼2.5 d

Steps 13–15 (Day 5), Passaging of expression cells: ∼2.5 d

Steps 16–20 (Day 7), Collection of the lentivirus-containing medium: ∼30 min

Steps 21–22 (Day 7), Infection of the expression cells: ∼2 d

Steps 23–24 (Day 9), Passaging of the transduced expression cells: ∼2.5 d

Steps 25–26 (Day 12), First antibiotic selection: ∼2 d

Steps 27–28 (Day 12), Passaging of backup cells: ∼3.5 d

Steps 29–30 (Day 14), Passaging of antibiotic-selected expression cells: ∼2.5 d

Steps 31–32 (Day 16), Second antibiotic selection: ∼2.5 d

Step 33–34 (Day 19), Passaging of polyclonal stable cell line: ∼3.5 d

Step 35 (Day 22), Expansion of polyclonal stable cell line, induction of protein expression and harvest: ∼2–3 weeks.

### Anticipated results

In a typical run, unconcentrated lentivirus-containing supernatant is sufficient to transduce HEK293-derived cells at modest MOI (∼0.5–1), yielding ∼40–60% infected cells. One to two cycles of antibiotic selection then eliminate non-transduced cells, resulting in a >95% enriched polyclonal population that regrows predictably, produces uniform expression, and is suitable for scale-up in either adherent or suspension formats. Dox-inducible control decouples selection and expansion from target protein expression, allowing induction only at the production stage, which is particularly advantageous for membrane proteins or proteins that burdensome or toxic to the host cells. Absolute production yields vary with target complexity, secretion or trafficking requirements, and culture format, but we routinely obtain milligrams to tens-of-milligrams per litre for both soluble and membrane proteins. These yields typically exceed those obtained with our original system ^1^, which we attribute to antibiotic enrichment preferentially retaining the highest-expressing cells within the production population.

Co-infection with two to three vectors results in increased early cell loss during multi-antibiotic selection, as only cells receiving all vectors survive; however, the surviving populations robustly co-express all components and are suitable for assembly of heteromeric complexes. In a typical advanced pHR-CMV-AB-TetO_2_ configuration, a TetR background maintained by blasticidin selection is combined with three lentiviral vectors carrying puromycin, zeocin, and hygromycin markers encoding subunits 1–3. Following multi-antibiotic co-selection, expression is synchronously induced with Dox, yielding a fully antibiotic-controlled workflow. Selected populations remain stable during routine passaging and can be cryopreserved and revived without loss of inducibility or expression level.

To demonstrate that antibiotic selection can recover uniform producer populations from sparse transduction, we infected HEK293T and HEK293S GnTI⁻ TetR cells with pHR-Puro-CMV-TetO_2_ lentiviruses encoding intracellular mNeonGreen (mNG), aiming for ∼5–15% mNG-positive cells prior to selection by adjusting the inoculum (Fig. 3b). After two selection cycles (24–72 h each), flow cytometry revealed reproducible enrichment to >90% mNG-positive with narrow, unimodal distributions, indicating efficient depletion of non-transduced cells and robust expression among selected cells (Fig. 4a,b). In HEK293S GnTI⁻ TetR, the enriched population exhibited low basal leakiness (∼1–5% mNG-positive without Dox), whereas addition of 1 µg/mL Dox produced a robust and uniform right-shift, demonstrating tight repression and synchronous induction (Fig. 4b). Together, these results demonstrate that antibiotic selection reliably converts sparsely transduced cell populations into near-uniform protein producers, with doxycycline providing precise temporal control over induction in TetR backgrounds.

**Fig. 4.**
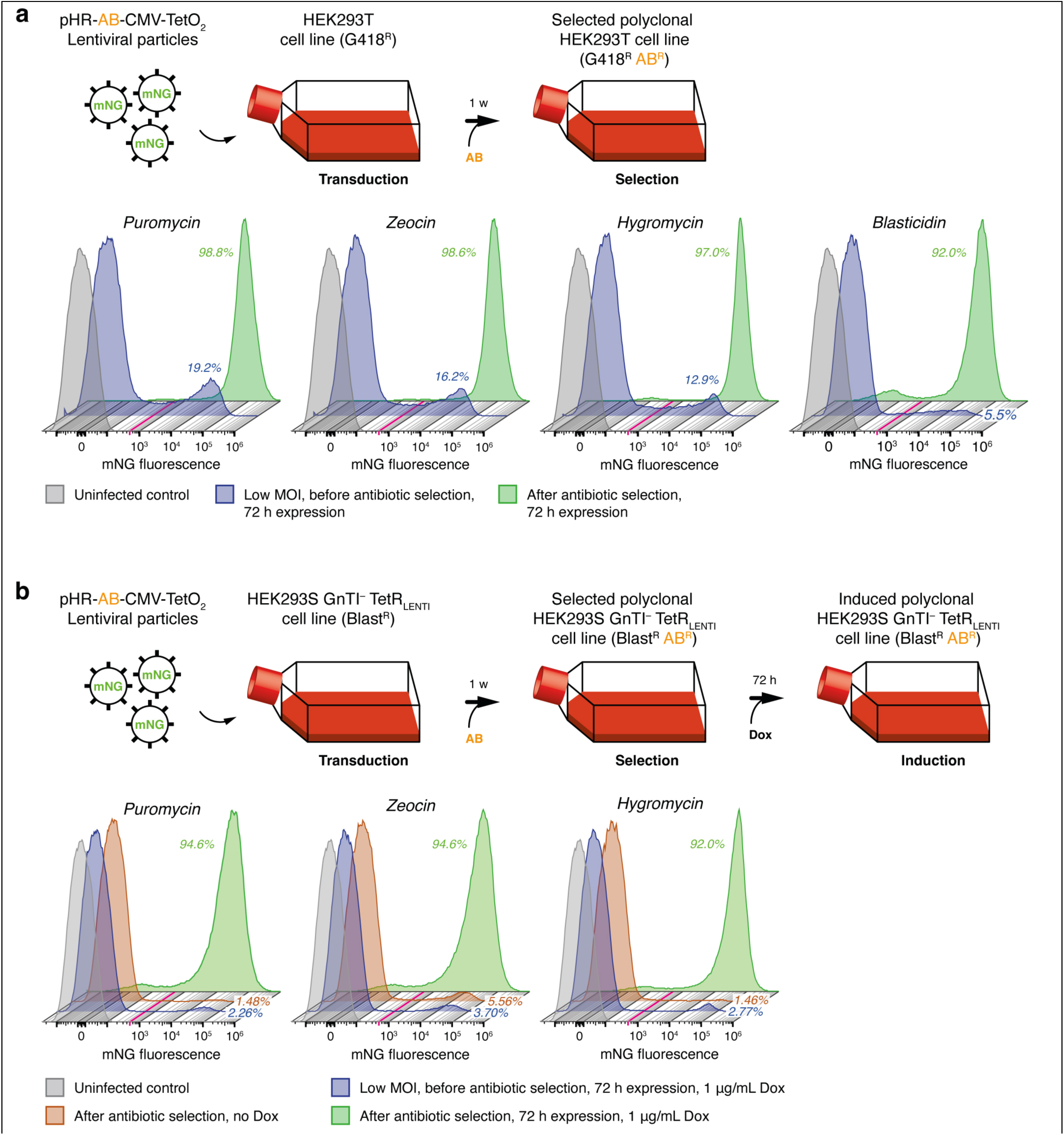
Antibiotic enrichment and Dox-inducible expression with pHR-AB-CMV-TetO_2_. **a**, HEK293T benchmark. Cells were infected at low MOI to yield a ∼5–15% mNG–positive cell pool before selection. Flow cytometry histograms show enrichment of the sparsely infected populations to >90% mNG-positive with a tight, unimodal distribution. Because HEK293T cells lack TetR, CMV-MIE-TetO_2_ drives constitutive expression. The magenta line marks the positivity threshold used for gating. **b**, HEK293S GnTI⁻ TetR benchmark. Low-MOI infection produced a ∼5 % mNG-positive cell pool that increased to >90% after two selection cycles. In the TetR background, basal expression in the enriched population was low-leak (∼1–5% positive without Dox); addition of Dox (1 µg/mL) produced a uniform and robust right-shift, indicating tight repression and synchronous induction. Gating as in (a). Abbreviations: AB, antibiotic; CMV, human cytomegalovirus; TetO₂, tandem Tet (tetracycline) operator sites; HEK293, human embryonic kidney 293; mNG, mNeonGreen; GnTI^−^, N-acetylglucosaminyltransferase I-deficient; TetR, Tet (tetracycline) Repressor protein; BlastR, blasticidin resistance; MOI, multiplicity of infection; Dox, doxycycline.

For the pHR-AIO-AB system in HEK293T cells, ∼5% mNG-positive cells were similarly targeted prior to selection. After one to two selection cycles (24–72 h), cultures were enriched to ∼80–90% mNG-positive with tight, unimodal distributions (Fig. 5a). Basal expression in the enriched AIO-AB populations was minimal (0–1% mNG-positive without Dox), whereas 1 µg/mL Dox induced a robust and uniform population-wide right-shift. Dox titration over a wide concentration range (0.06–3000 ng/µL) yielded sigmoidal dose–response curves, with half-maximal activation at 0.37 ng/mL (0.33–0.41) for the fraction of mNG-positive cells and at 1.59 ng/mL (1.41–1.78) for median fluorescence intensity (MFI), indicating the high sensitivity of the pHR-AIO-AB configuration to Dox, and demonstrating that Dox dose can be used to tune expression levels while maintaining tight regulation and population homogeneity (Fig. 5b).

**Fig. 5.**
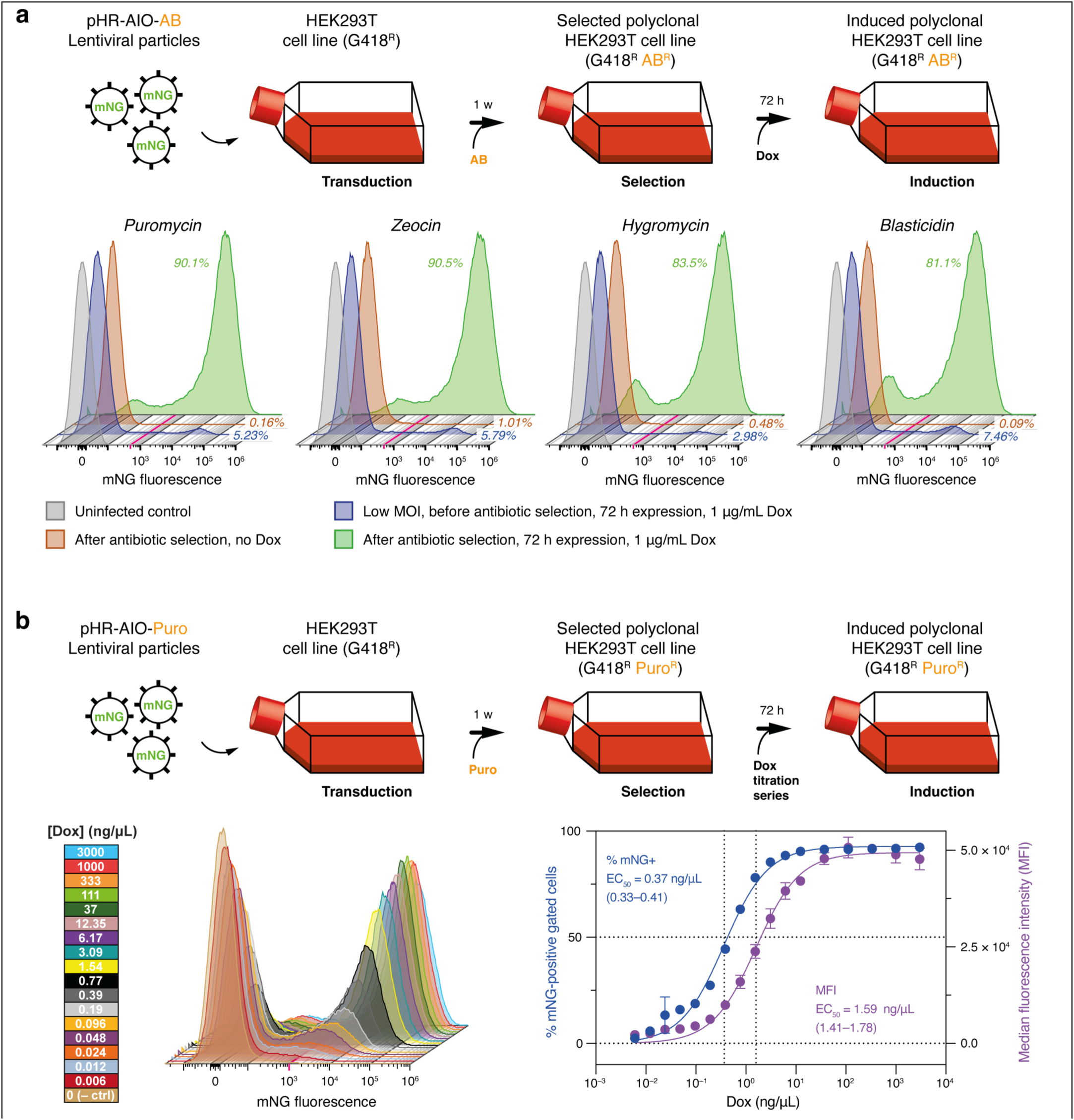
Antibiotic enrichment and Dox-inducible expression with pHR-AIO-AB. **a**, HEK293T cells were infected at low MOI to yield a ∼5% mNG–positive cell pool before selection. Flow cytometry histograms reveal enrichment of the sparsely infected populations to >80% mNG-positive with a tight, unimodal distribution. Basal expression in the enriched population was low-leak (∼0–1% positive without Dox); addition of Dox (1 µg/mL) produced a uniform and robust right-shift, indicating tight repression and synchronous induction. The magenta line marks the positivity threshold used for gating. **b**, Doxycycline titration of pHR-AIO-AB-mediated expression in HEK293T cells. HEK293T cells were transduced with pHR-AIO-Puro lentiviral particles encoding mNG, enriched by puromycin selection, and subsequently induced using a doxycycline (Dox) titration series. Bottom left, overlaid flow-cytometry histograms showing mNG fluorescence distributions across increasing Dox concentrations (0–3000 ng/mL), illustrating a progressive and uniform right-shift of the population with minimal basal expression in the absence of Dox. Bottom right, dose–response curves plotting the fraction of mNG-positive cells (blue, left y-axis) and the median fluorescence intensity (MFI; purple, right y-axis) as a function of Dox concentration. Sigmoidal fits reveal half-maximal activation (EC_50_) at 0.37 ng/mL (0.33–0.41) for the fraction of mNG-positive cells and at 1.59 ng/mL (1.41–1.78) for MFI, demonstrating high sensitivity and tunable induction across the population. Abbreviations: AIO, all-in-one; mNG, mNeonGreen; Dox, doxycycline; MOI, multiplicity of infection; MFI, median fluorescence intensity. Abbreviations: AIO, all-in-one; AB, antibiotic; HEK293, human embryonic kidney 293; mNG, mNeonGreen; MOI, multiplicity of infection; Dox, doxycycline; Puro, puromycin; MFI, median fluorescence intensity.

## Author contributions

J.E. designed and constructed the pHR-CMV-AB-TetO_2_ and pHR-AIO-AB plasmid suites. E.B., A.N., A.D. and J.E. established and optimised the protocol. E.B. and J.E. performed the flow cytometry. E.B. and J.E. wrote the paper, with contributions from all authors.

## Acknowledgments

This work was supported by the European Research Council (ERC) Starting Grant 850820 (SynLink) to J.E., and the Région Nouvelle-Aquitaine grant 3428720 to J.E. We thank Justine Charpentier (IINS, CNRS UMR 5297) for support with molecular biology. pMD2.G was a gift from Didier Trono (Addgene plasmid # 12259; http://n2t.net/addgene:12259; RRID:Addgene_12259). psPAX2 was a gift from Didier Trono (Addgene plasmid # 12260; http://n2t.net/addgene:12260; RRID:Addgene_12260).

## Competing interests

The authors declare no competing interests.

## Data availability

All pHR-AB-CMV-TetO_2_ and pHR-AIO-AB transfer plasmids described in this protocol are available via Addgene (https://www.addgene.org/) and are distributed under a Uniform Biological Material Transfer Agreement (UBMTA). The engineered HEK293-derived cell lines (including HEK293T Lenti-X ΔPKR, HEK293T TetR_LENTI_, and HEK293S GnTI^−^ TetR_LENTI_) are available upon request from the corresponding author. No additional restrictions apply beyond standard material transfer requirements for plasmids (UBMTA via Addgene) and distribution of cell lines by direct request.

## Notes

### Competing Interest Statement

The authors have declared no competing interest.

